# *Mycobacterium ulcerans* in Mosquitoes and March flies captured from endemic areas of Northern Queensland, Australia

**DOI:** 10.1101/390997

**Authors:** Avishek Singh, William John Hannan McBride, Brenda Govan, Mark Pearson, Scott A. Ritchie

## Abstract

*Mycobacterium ulcerans* is the causative agent of Buruli ulcer (BU). This nontuberculous mycobacterial infection has been reported in over 33 countries worldwide. In Australia, the majority of cases of BU have been recorded in coastal Victoria and the Mossman-Daintree areas of north Queensland. Mosquitoes have been postulated as a vector of *M. ulcerans* in Victoria, however the specific mode of transmission of this disease is still far from being well understood. In the current study, we trapped and analysed 16,900 (allocated to 845 pools) mosquitoes and 296 March flies from the endemic areas of north Queensland to examine for the presence of *M. ulcerans* DNA by polymerase chain reaction. Seven of 845 pools of mosquitoes were positive on screening using the IS2404 PCR target but only one pool was positive for presence of *M. ulcerans* after confirmatory testing. None of the March fly samples were positive for the presence of *M. ulcerans*. *M. ulcerans* was detected on proboscises of deliberately exposed mosquitoes.

**Author Summary:** The causative agent of Buruli ulcer is Mycobacterium ulcerans. This destructive skin disease is characterized by extensive and painless necrosis of skin and underlying tissues usually on extremities of body due to production of toxin named mycolactone. The disease is prevalent in Africa and coastal Australia. The exact mode of transmission and potential environmental reservoir for the pathogen still remain obscure. Aquatic and biting insects have been identified as important niche in transmission and maintenance of pathogen in the environment. In this study we screened mosquitoes and march flies captured from endemic areas of northern Queensland for the presence of *M. ulcerans.* In addition, we conducted artificial blood feeding experiment to identify the role of mosquitoes in transmission of this pathogen. We found one pool of mosquito out of 845 pools positive for *M. ulcerans* and none of the March fly samples were positive. This could indicate a low burden of the bacteria in the environment coinciding with a comparatively low number of human cases of *M. ulcerans* infection seen during the trapping period of the study. Evidence to support mechanical transmission via mosquito proboscises was found.

## Introduction

Buruli ulcer (BU), also known regionally as Daintree ulcer in north Queensland, Australia or Bairnsdale ulcer in Victoria, Australia, is an emerging disease of skin and underlying tissue, with a potential to lead to permanent disability, particularly if treatment is inadequate or delayed. The causative agent of this disease, *M. ulcerans* secretes a polyketide exotoxin, mycolactone, the production of which requires expression of a series of contiguous genes on the large pMUM001 plasmid. This exotoxin is the main virulence determinant of the bacteria (1). The outbreaks of BU have been consistently linked with wetland or coastal regions (2). Environmental samples such as water, aquatic plants, soil at endemic areas has been found PCR-positive for *M. ulcerans* DNA (3, 4). Insects such as mosquitoes and aquatic bugs has been proposed as a vital ecological niche for the maintenance of pathogen in environment (5, 6). The detection of *M. ulcerans* DNA in insects does not prove their ability to transmit *M. ulcerans* but could indicate potential to act as either biological or mechanical vector. A study conducted by Marsollier and his colleagues provided evidence of the presence of *M. ulcerans* in the salivary gland of wild caught Naucoridae (aquatic bug). They successfully isolated the pathogen by culture from the salivary glands of aquatic bugs and suggested aquatic insects as having an important ecological niche in the maintenance of the organism in the environment. They were also able to demonstrate transmission to mice in a laboratory environment (6). Similarly, a study conducted by Wallace *et al*. provided evidence of the ability of mosquitoes to act as a mechanical vector of *M. ulcerans* (7). Studies conducted in endemic areas of Africa suggest that conducting farming activities close to rivers (8) and swimming in rivers located in endemic areas (9) are risk factors for exposure to *M. ulcerans.*

In Australia, foci of BU infection have been found in tropical Far North Queensland (10, 11), the Capricorn Coast region of central Queensland (10), the Northern Territory (12) and temperate coastal Victoria (5). Victorian researchers detected the presence of *M. ulcerans* in five different species of mosquito during a BU outbreak in an endemic area of Victoria, Australia. They demonstrated the absence of *M. ulcerans* in a neighbouring area, where BU did not occur (5). Together, the evidence was proposed to support a link with mosquitoes in the ecology of BU in Victoria (5, 13). More recently, there was a report of the presence *M. ulcerans* in a single mosquito out of two pools collected in the BU endemic region of north Queensland. The isolated detection of *M. ulcerans* in a tropical endemic region in Australia highlighted a need to examine a larger sample size to gauge the significance of the role of mosquito in ecology of BU in Northern Queensland (14). An additional hypothesis put forward by the local population (including people with a history of BU) was that March flies (Tabanidae) might have a role in transmission (Villager *et al*, unpublished manuscript). We therefore aimed, in this study to capture and screen mosquitoes and March flies for the presence of *M. ulcerans* DNA in the BU endemic area of Northern Queensland. In addition, we conducted a mosquito artificial blood feeding experiment to demonstrate an in vitro basis for mechanical transmission of *M. ulcerans* by blood fed mosquitoes.

## Material and Methodology

Selection of the study site was based on GIS mapping of human cases of BU in Northern Queensland (15). We divided the endemic area of northern Queensland into three regions: Region-1: extending from Miallo to lower Daintree including Wonga/Wonga Beach area, Region-2: Forest Creek area and Region-3: Upper Daintree area for ease of sampling and analysis (Fig. 1).

**Fig. 1:**
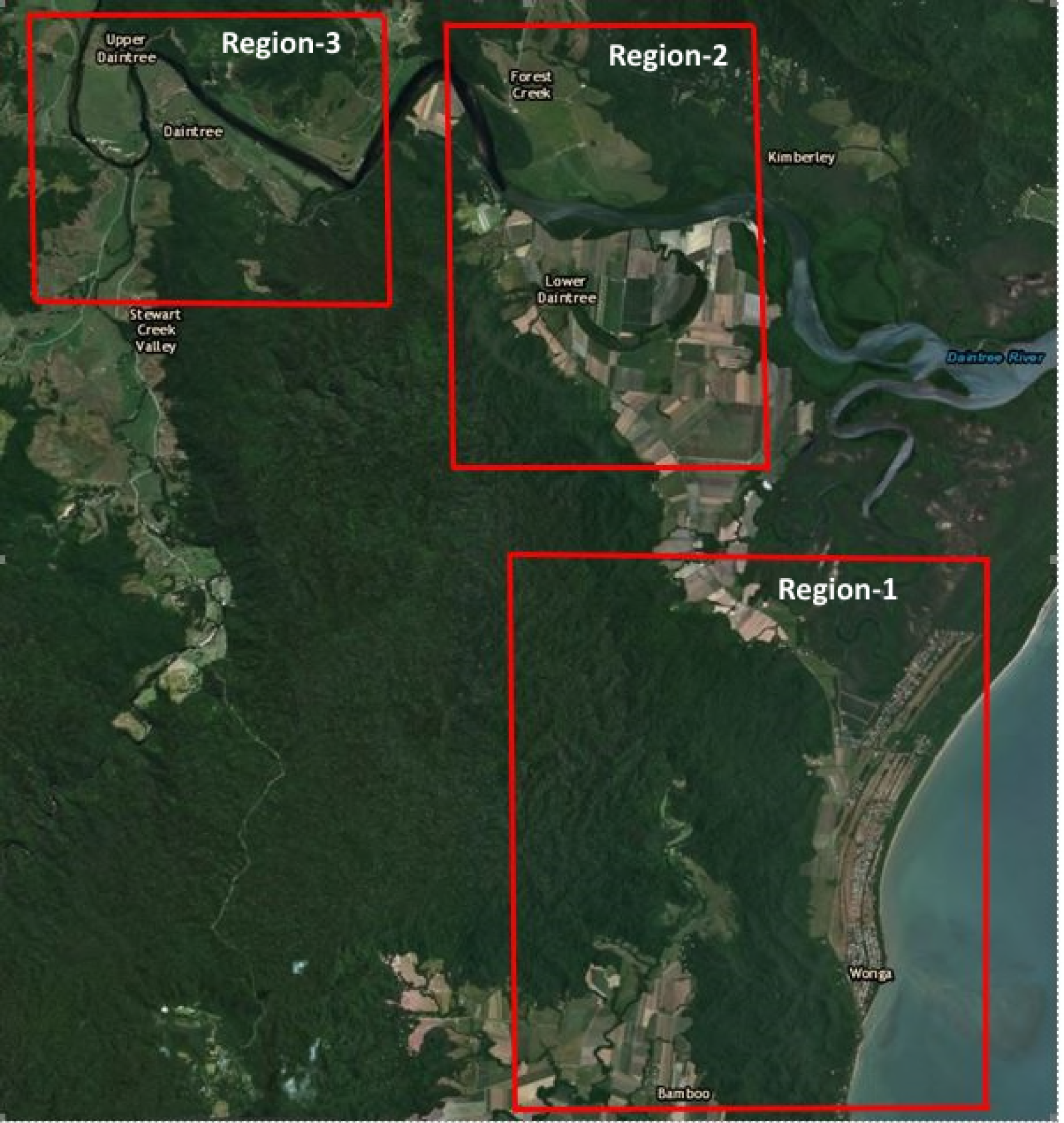
BU endemic areas of Northern Queensland, Australia and Mosquito trapping regions. This figure was created using base layer obtained from https://landsatlook.usgs.gov/.

### Trapping of Mosquitoes

Mosquitoes were captured using a model 512 "CDC miniature light trap" (John W. Hock Company, Gainesville Florida USA) baited with 1 kg of dry ice as the source of CO_2_. This trap is the most reliable, efficient and portable device for trapping mosquitoes and sand flies (16). This trap consists of an electric light and fan just over the collection container and is operated by a 12V battery. A two liter insulated container was used to hold dry ice and a pipe was attached to release CO_2_ over the trap to attract mosquitoes (Fig. 2). Thirty overnight trapping sessions were conducted starting from September 2016 through to February 2018, with at least 4 CDC traps placed within a 1 kilometer radius of each-other. Of the 30 trapping sessions, 14 were conducted at eight different sites within region-1, nine at six different sites within region-2 and seven at five different sites of region-3 (Fig. 1). Traps were placed at different sites after obtaining permission to access properties from the owners and selection of sites were based on history of BU cases in humans in nearby households. Geographical Information System (GIS) coordinates of each trap was recorded. On each occasion, traps were set before dusk and checked for mosquitoes after dawn the next morning. After each occasion of trapping, catches were transported to the Mosquito Research Facility, Australian Institute of Tropical Health and Medicine (AITHM), James Cook University, Cairns, Australia where they were counted, sorted and pooled by genus, with each pool containing ≤ 20 mosquitoes of same genus and collected from the same site.

**Fig. 2.**
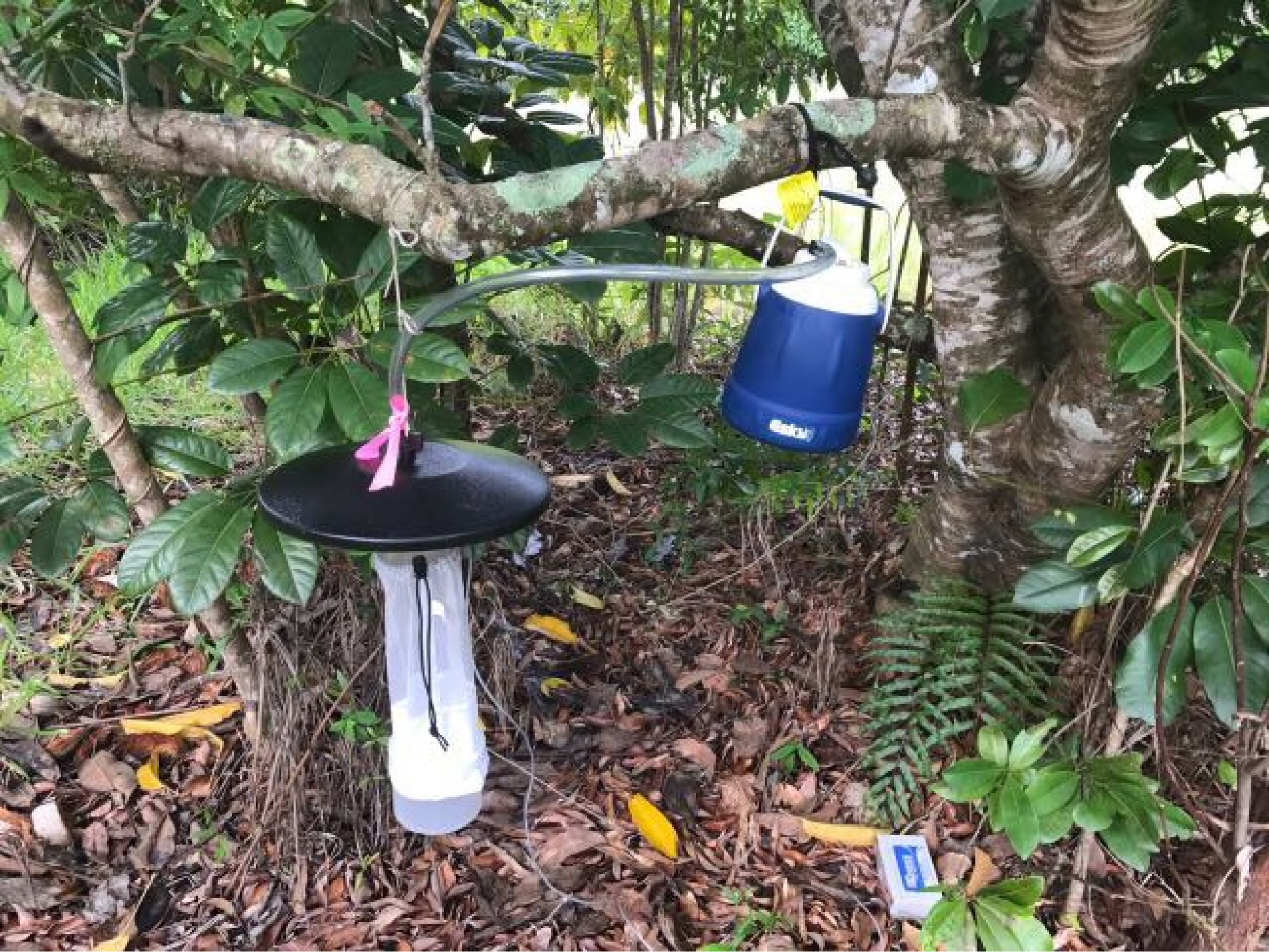
CDC miniature light trap baited with dry ice.

### Trapping of March Flies

Several attempts were made to trap march flies from endemic areas with an investigator wearing dark clothes to attract them, or with the use of an insect net sprayed with insecticide. These attempts occurred from February 2016 through September 2016. The yield from these attempts were very low. A request was made to residents of region-1 through the local State School to collect march flies. This effort was successful and large numbers of March flies of genus *Tabanus* were collected by the local community. The addresses of properties from which March flies were collected were recorded. Sampling of March flies was restricted to region-1.

## Screening of Mosquitoes and March Flies for MU DNA by PCR

DNA was extracted from each pools of × 20 mosquitoes of the same genus by using the FastPrep Instrument (MP Biomedicals, Solon, OH, USA) as per manufacturer’s instruction with FastDNA Kit (MP Biomedicals). Using the same instrument, DNA from individual March fly was extracted with FastDNA Spin Kit (MP Biomedicals). Extracted DNA was stored at −20 °C. The extracted DNA samples were screened for the presence of *M. ulcerans* DNA by using a semi-quantitative real-time PCR adapted from a method for the detection of *M. ulcerans* DNA from environmental samples (17). To rule-out the possibility of contamination, three negative controls (double deionized water, MilliQ) and three positive controls (purified *M. ulcerans* DNA obtained from Victorian Infectious Disease Reference Laboratory) were used during RT-PCR assay run. All of the extracted DNA samples were initially screened for the *M. ulcerans* insertion sequence element IS 2404. Samples positive for IS 2404 were re-analyzed by a second real-time PCR for the detection of two additional regions in the genome of *M. ulcerans*: IS 2606 and ketoreductase B domain (KR). This screening process has been validated by Fyfe *et al*. to differentiate *M. ulcerans* from other mycolactone producing mycobacteria (MPM) (17). They suggested that the difference in real-time PCR cycle thresholds (Ct) between IS 2606 and IS 2404 (ΔCt [IS 2606 - IS 2404]) allows for the differentiation of *M. ulcerans* from other MPM that contain IS 2404 but which have fewer copy numbers of IS 2606. The detection of all three targets (IS 2404, IS 2606 and KR) with an expected ×Ct values (< 7) confirms the presence of *M. ulcerans* DNA in the sample. (17, 18).

## Mosquito artificial blood feeding experiment

An isolate of *M. ulcerans* obtained from the Mycobacterium Reference Laboratory at the Royal Brisbane Hospital was used for this experiment. *M. ulcerans* was confirmed via PCR analysis. Our laboratory was not equipped with facilities to safely conduct transmission experiments, so isolates of *M. ulcerans* were subjected to UV light to kill the pathogen. To confirm the sterilty of the isolates, an aliquot was sub-cultured onto Lowenstein-Jensen (LJ) slants and liquid Middlebrook 7H9 media supplemented with 10% oleic acid-albumin-dextrose enrichment (OADC). Inoculated media was kept at 31°C in 25cm2 tissue culture flasks and observed for 8 weeks for growth. No growth was observed confirming the absence of live bacteria. The killed isolates were subjected to RT-PCR targeting IS 2404, IS 2606 and KR as described above. RT-PCR analysis confirmed the presence of *M. ulcerans* DNA.

We used an artificial blood feeding method (simple membrane method) described by Finlayson *et al*. with some modification in this study (Finlayson, Saingamsook, & Somboon, 2015). This procedure is a simple and affordable alternative for direct host feeding (DHF). The method involves pouring warmed defibrinated sheep blood (Applied Biological Products Management-Australia) into the indented base on the underside of a plastic container and then covering it with a stretched collagen membrane secured by a rubber band. The container is then turned up, filled with warm water and covered by a lid. The feeder is then placed on the mesh side of the cage, allowing the mosquitoes to pierce the collagen membrane to access the blood.

The experiment was conducted using wild type *Aedes aegypti* hatched and reared in the same batch, sorted as pupae into four cages containing 30 female mosquitoes in each. One out of four cages was used as control (Cage-D) where only defibrinated sheep blood was used as feed and in remaining three cages (Cage-A, B and C) defibrinated sheep blood mixed with killed *M. ulcerans* isolates were used. All four cages were exposed to blood for 2 hours. Fully blood fed mosquitoes from each cage were aspirated separately and knocked down by freezing. Pool of mosquitoes from cage A, and B were dissected separating the head, abdomen and legs of each insects by sterile fine forceps to avoid contamination during dissections. DNA from the head, abdomen and legs (pooled separately) from the mosquitoes from cage A and B and whole mosquitoes from cage C and D were extracted using FastPrep Instrument (MP Biomedicals, Solon, OH, USA) as per manufacturer’s instruction with FastDNA Kit (MP Biomedicals). All the extracted DNA were initially screened for IS 2404 and IS 2404-positive samples were re-analyzed for IS 2606 and KR with RT-PCR assay as described above.

## Results

### Screening of Mosquitoes

A total of 16,900 mosquitoes were captured over the course of the study from 30 occasions of trapping at three different regions of northern Queensland. Total mosquitoes captured from region-1, region-2 and region-3 were 7880, 5100, and 3920, respectively. The majority of captured mosquitos belonged to the *Verrallina* genus (specifically *Verrallina lineata*) 82%, followed by *Coquillettida* (9%) and *Mansonia* (3%). The remaining 6% consisting seven other genera that were classified as "other" for screening. See figure 3 below.

**Fig. 3.**
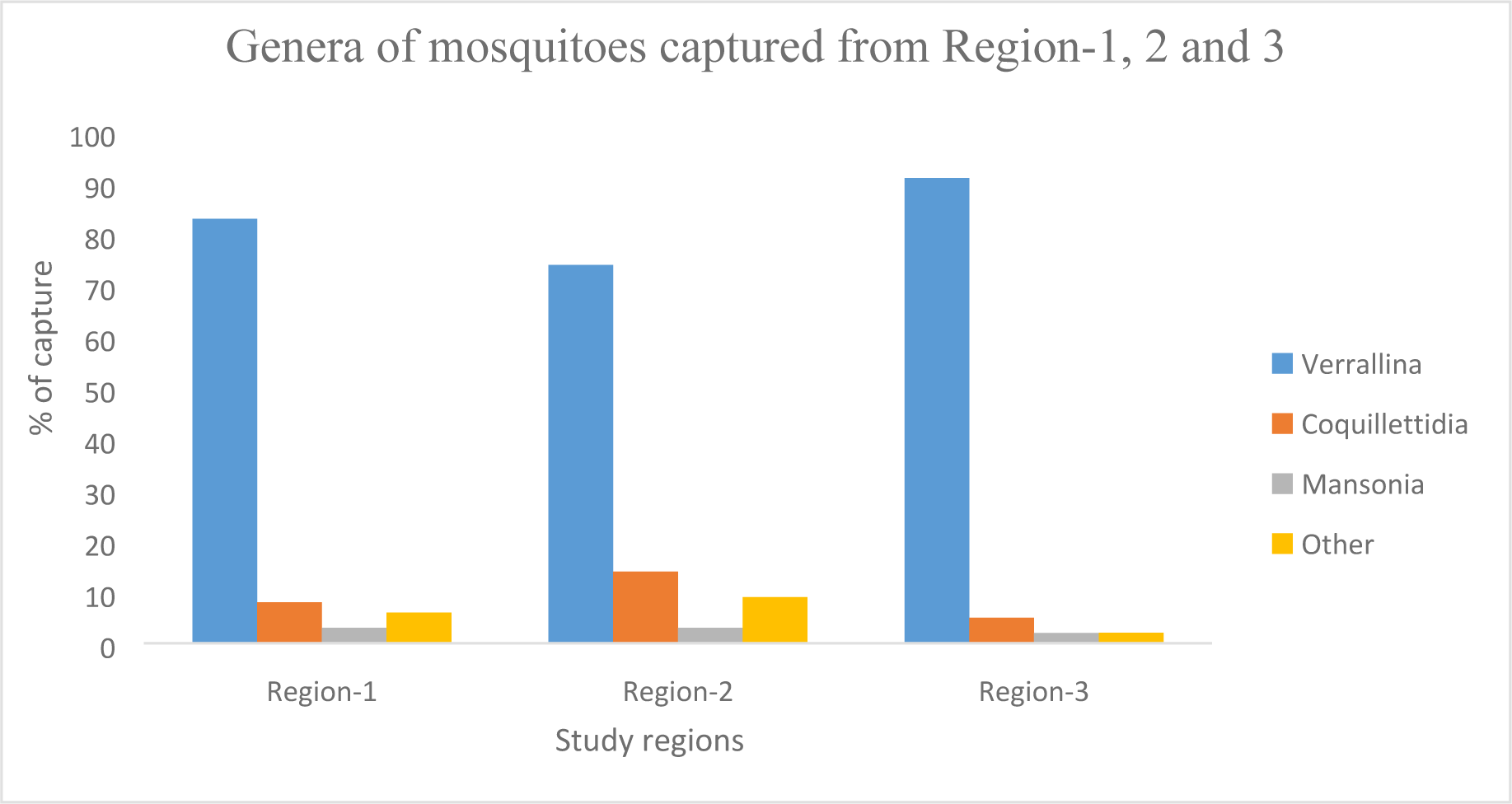
Genera of mosquitoes captured from three different regions: Region-1 comprising 83% of *Verrallina sp.*, 8% of *Coquillettida sp.*, 3% of *Mansonia sp*. and 6% of others; Region-2 comprising 74% of *Verrallina sp.,* 14% of *Coquillettida sp.*, 3% of *Mansonia sp.* and 9% of others and Region-3 comprising 91% of *Verrallina sp.,* 5% of *Coquillettida sp.*, 2% of *Mansonia sp.* and 2% of others of total catches.

Of total 16,900 mosquitoes screened (845 pools), seven pools were positive for IS 2404. Three of those seven pools were *Verrallina sp.* from region-1, two pools were *Coquillettida sp.* one each from capture region-1 and 3 and the remaining two pools were *Mansonia sp.* from region-1. Of the seven pools positive for IS 2404, one pool was positive for *M. ulcerans* using the confirmatory PCR assays. This single positive pool of mosquitoes was comprised of genus *Verrallina sp.* and linked to region-1. Thirty pools of mosquitoes which were negative for IS 2404 were tested for IS 2606 and KR. None of them were positive for these probes signifying the dependent nature of existence of IS 2606 and KR with IS 2404. Similar findings were reported during the Victorian outbreak (5).

### Screening of March flies

DNA extracts of 296 March flies were screened for IS 2404. None of the samples were positive for this probe. Twenty-four randomly selected IS 2404 negative samples were tested for IS 2606 and KR and none were positive.

### Mosquito artificial blood feeding experiment

There were a total of seven samples: 2 pools of heads, 2 pools of abdomens, 2 pools of legs (from cage A and B) and 1 pool of whole mosquitoes (Cage C). DNA extracted from pools of heads and abdomens of mosquitoes from cage A and B and the pool of whole mosquitoes from Cage C were positive for IS-2404. Confirmatory assays targeting IS 2606 and KR revealed that three samples: DNA extract from the heads of mosquitoes from cage A and B and the pool of mosquitoes from cage C were positive for *M. ulcerans* DNA. Controls (a pool of mosquitoes from cage D) were negative for all three targets: IS 2404, IS 2606 and KR.

## Discussion

Mosquitos serve as important biological vectors for a variety of pathogens. The movement of pathogens from the gastro-intestinal tract after ingestion to the salivary glands for subsequent transmission is well documented for many diseases. However, this phenomenon has not been demonstrated for *M. ulcerans*. A study conducted by Wallace and colleagues (2010) provided evidence on the maintenance of *M. ulcerans* throughout larval development without further passage of the organisms into pupa or adult mosquitoes (19). They concluded that mosquitoes were an unlikely biological vector of *M. ulcerans*. Wallace *et al* (2017) subsequently provided evidence of mechanical transmission of *M. ulcerans* via anthropogenic skin puncture or mosquito bites (7). Detection of *M. ulcerans* from pools of head of mosquitoes in our mosquito artificial blood feeding experiments indicate a potential for mosquitoes act as an agent for mechanical transmission of *M. ulcerans*. However, our mosquito artificial blood feeding experiment had some limitations. We were not able to conduct experiments to verify whether *M. ulcerans* positive mosquitoes transmit pathogen to healthy animal or not. Only proboscises of mosquitoes were in direct contact with blood containing *M. ulcerans* DNA. *M. ulcerans* was only detected from the abdomen of mosquitoes using the IS2404 target. This might have been due to insufficient amount of DNA to identify IS 2606 and KR.

For mechanical transmission, insect vectors such as mosquitoes must acquire the pathogen either from the environment or an infected host. For this to occur efficiently, the organism must be abundantly present in the environment. A survey in Victoria, Australia has confirmed a strong correlation between mosquitoes found to test positive for carrying *M. ulcerans* and the number of human cases of BU occurring (5). The group found a significantly higher number of mosquitoes screened positive for *M. ulcerans* during an intense outbreak of BU in endemic areas, in comparison to areas with a lower incidence of human cases.

The number of human cases of BU has decreased in Northern Queensland, Australia since the largest recorded outbreak in 2011 (> 60 cases). The majority of the cases during the 2011 outbreak were from Wonga and the Wonga beach area, referred as region-1 in the study by Steffen and Freeborn (2018) (20). Out of 394 pools collected, only one from region-1 was positive for *M. ulcerans* in this study. Interestingly, mosquitoes of this positive pool were trapped in the backyard of a property in Wonga Beach area (region-1) where two human cases of BU were confirmed in 2017. All other pools of mosquitoes and march flies collected from that properties negative for *M. ulcerans*.

In a separate study conducted in Northern Queensland, Australia, one pool of mosquitoes was found positive for *M. ulcerans* out of two pools collected in total (14). However, it must be noted that this study was conducted soon after 2011 which raises the possibility that sampling should occur as close as possible in time to when transmission is thought to be occurring.

*M. ulcerans* is an environmental pathogen and detection of *M. ulcerans* positive mosquitoes may only be an indicator for the presence of the organism in the environment. A significant decrease in human cases of BU in Northern Queensland in recent years could be due to a lower load of bacteria in the environment. This may explain the low detection of *M. ulcerans* positive mosquitoes and March fly populations in the study sites. However, the detection of *M. ulcerans* even in a single pool of mosquitoes from the endemic areas of Northern Queensland is significant, as it corroborates findings in Victoria where five different species of mosquitoes captured from BU-endemic regions during human outbreaks were positive for *M. ulcerans*.

Our detection of *M. ulcerans* in mosquitoes in Northern Queensland does support the earlier report from Victoria in Australia (5). The Victorian study provides evidence for high detection rates of *M. ulcerans* positive mosquitoes if captured during peak times of outbreaks. Our study found that it is less likely to find *M. ulcerans* positive mosquitoes if they are trapped from areas where human incidence of BU is currently low. We hypothesize that mosquitoes and perhaps other biting insects, such as March flies may have a significant role in the ecology and transmission of *M. ulcerans* in endemic areas during outbreaks and that the level of detection of *M. ulcerans* positive mosquitoes in the environment could be an indicator for disease outbreaks.

## Acknowledgements

We thank laboratory staff of Mosquito Research Facility, AITHM for their technical advice trapping of mosquitoes and designing blood feeding experiments and Janet Fyfe from Mycobacterium Reference Laboratory at VIDRL for her technical advice in analyzing samples. We are grateful to Wonga Beach State School for their assistance in collection of March flies. We thank Hendrik Weimar and local community for their continuous support and assistance in arranging access to the sites for setting traps.

